# Increased mTOR signaling, impaired autophagic flux and cell-to-cell viral transmission are hallmarks of SARS-CoV-2 infection

**DOI:** 10.1101/2021.10.13.464225

**Authors:** Grazielle Celeste Maktura, Thomaz Luscher Dias, Érika Pereira Zambalde, Bianca Brenha, Mariene R. Amorim, Luana Nunes Santos, Lucas Buscaratti, João Gabriel de Angeli Elston, Cynthia Mara, Mariana Camargo Silva Mancini, Isadora Carolina Betim Pavan, Daniel A. Toledo-Teixeira, Karina Bispo-dos-Santos, Pierina L. Parise, Stefanie Primon Muraro, Gabriela Fabiano de Souza, Ana Paula Morelli, Luiz Guilherme Salvino da Silva, Ícaro Maia Santos de Castro, Guilherme O. Barbosa, Raissa G. Ludwig, Thiago L. Knittel, Tatiana D. Saccon, Marcelo A. Mori, Fabiana Granja, Hernandes F. Carvalho, Luis Lamberti Pinto da Silva, Helder I. Nakaya, Jose Luiz Proenca-Modena, Fernando Moreira Simabuco, Henrique Marques-Souza

**Author notes:** These authors contributed equally to this work (Grazielle Celeste Maktura, Thomaz Luscher Dias, Érika Pereira Zambalde, Bianca Brenha, Mariene R. Amorim).

## Abstract

The COVID-19 disease caued by the Severe Acute Respiratory Syndrome Coronavirus 2 (SARS-CoV-2) has two characteristics that distinguish it from other viral infections. It affects more severely people with pre-existing comorbidities and viral load peaks prior to the onset of the symptoms. Investigating factors that could contribute to these characteristics, we found increased mTOR signaling and suppressed genes related to autophagy, lysosome, and vesicle fusion in Vero E6 cells infected with SARS-CoV-2. Transcriptomic data mining of bronchoalveolar epithelial cells from severe COVID-19 patients revealed that COVID-19 severity is associated with increased expression of genes related to mTOR signaling and decreased expression of genes related to autophagy, lysosome function, and vesicle fusion. SARS-CoV-2 infection in Vero E6 cells also resulted in virus retention inside the cells and trafficking of virus-bearing vesicles between neighboring cells. Our findings support a scenario where SARS-CoV-2 benefits from compromised autophagic flux and inhibited exocytosis in individuals with chronic hyperactivation of mTOR signaling, which might relate to undetectable proliferation and evasion of the immune system.

## 1. Introduction

The new coronavirus disease (COVID-19) caused by Severe Acute Respiratory Syndrome Coronavirus 2 (SARS-CoV-2) became a pandemic, severely affecting people with pre-existing comorbidities, such as hypertension, diabetes, obesity, and heart disease [1,2]. Increased signaling through the mammalian/mechanistic Target of Rapamycin (mTOR) signaling pathway is a common characteristic in all comorbidities associated with a higher risk of mortality by COVID-19 [3–8]. In healthy individuals, mTOR signaling is responsible for maintaining a balance between protein synthesis, autophagy, and nutrient storage processes. This balance is crucial for the cell since its dysregulation leads to cancer, obesity, and diabetes [9]. Chronic mTOR hyperactivation is found in several diseases [10]. The process of autophagy, from autophagosome formation until lysosomal degradation, depends on the inactivation of mTOR signaling [11,12]. Interestingly, an interactomics study identified mTOR signaling as a pathway possibly modulated by SARS-CoV-2 infection [13]. Also, recent findings showed that SARS-CoV-2 infected cells have compromised autophagic flux and increased virus replication [14,15]. Therefore, it is tempting to predict that the balance between mTOR and autophagy could link comorbidities to severe COVID-19 symptoms.

Differently from the Middle East Respiratory Syndrome Coronavirus (MERS-CoV) and the Severe Acute Respiratory Syndrome Coronavirus (SARS-CoV), SARS-CoV-2 viral load peaks before the onset of the symptoms and remains elevated for up to three weeks [16,17]. It is still unknown the reason why SARS-CoV-2 can proliferate and remain undetectable by the immune system during the early stages of the disease. Interestingly, high mTOR and low lysosomal activation correlate with reduced exocytosis [6,18], suggesting a possible link between mTOR, autophagy, and virus release from infected cells.

In this report, we analyzed molecular pathways of SARS-CoV-2 infection in Vero E6 cells and COVID-19 patients to evaluate the balance between the mTOR signaling pathway and the process of autophagy and their correlation with viral release and transmission by cell-to-cell connections, a phenomenon usually associated with viral persistence and immune evasion. Our findings reveal that SARS-CoV-2 infection in Vero E6 cells and severe COVID-19 patients occur in conditions of increased mTOR activity and impaired autophagic flux that could prevent the viral release and promote cell-to-cell viral transmission. These data may help to explain why the SARS-CoV-2 load peak occurs before COVID-19 symptoms and the exacerbated activation of the immune response seen in severely affected COVID-19 patients.

## 2. Materials and Methods

### Vero E6

Vero E6 (African green monkey, *Cercopithicus aethiops*, kidney) cells were cultured in Dulbecco’s modified Eagle medium (DMEM; Sigma-Aldrich, USA) supplemented with 10% heat-inactivated fetal bovine serum (FBS; Sigma-Aldrich, USA) and 1% Penicillin-Streptomycin 100 U/mL and 100 μg/mL (Sigma-Aldrich, USA), and incubated in 5 % carbon dioxide atmosphere at 37 °C.

### Viral infection

An aliquot of SARS-CoV-2 SP02.2020 (GenBank accession number MT126808) isolate was kindly donated by Professor Edison Luiz Durigon of the University of São Paulo, São Paulo [19]. Vero E6 cells were used for virus propagation in the Biosafety Level 3 Laboratory (BSL-3) of the Laboratory of Emerging Viruses. Viral infections were performed in Vero cells seeded in 24 wells plates (5 × 10^5^ cells/well) for the experiments with treatments and immunofluorescence assays, and 6 wells plates (1 × 10^6^ cells/well) for Western blots. A multiplicity of infection (MOI) of 1 was used for all experiments.

### Plaque-forming units’ assay (PFU)

PFU assay was performed to access viral titer in the supernatant of infected cells and experimental samples as previously described, with modifications [19,20]. Briefly, 24 wells plates were seeded with Vero E6 cells at a quantity of 5 × 10^5^ cells/well. Six 10-fold dilutions of the samples were performed and transferred as duplicates to the wells. Positive and negative control of SARS-CoV-2 was included. The plates were incubated for 1 h for adsorption in a rocking platform at room temperature. Then, a semisolid medium, containing 1 % carboxymethyl cellulose (CMC) (Sigma-Aldrich, USA) in DMEM, 2 % FBS, and 1 % of penicillin-streptomycin, was added to the wells. The plates were incubated at 37 °C, 5 % of CO_2_ atmosphere for 48 h. The semisolid medium was then removed, cells were fixed with paraformaldehyde 8 % for 30 minutes, washed with distilled water, and stained with methylene blue (Sigma-Aldrich, USA).

To access the production of viable viral particles inside Vero cells in the experiments, the supernatants were removed from the wells, cells were washed once with PBS 1X, and the monolayer was collected and frozen with liquid nitrogen, followed by three series of freeze-and-thaw in liquid nitrogen and 37 °C water bath. These samples were titrated by PFU.

### Bafilomycin treatment

For the bafilomycin experiment, cells were cultured in 24-well plates as described above, followed by washing with phosphate-buffered saline (PBS) and treated with 10 nM of bafilomycin in DMEM 10% FBS for 3 hours. Treatment was withdrawn two hours prior to infection and 24 hpi [21,22] cells and supernatant samples were titrated by PFU as previously described.

### Probe synthesis

The first step to generate the probe was to amplify the RdRp gene target from SARS-CoV-2 cDNA using GoTaq® DNA Polymerase PCR kit (Promega) and the following oligonucleotide primers RdRp Fwd 5’-AACACGCAAATTAATGCCTGTCTG 3’ and RpRd Rev 5’ GTAACAGCATCAGGTGAAGAAACA 3’. The polymerase chain reaction 1 (PCR 1) was carried out in a total volume of 50 μL containing 10 μL 5X GoTaq reaction buffer, 2 μL MgCl2, 1 μL dNTP mix (10 mM), 1 μL of each RdRp primer (10 mM), 1 μL cDNA (1:10 dilution), 0.25 μL GoTaq® DNA Polymerase (5 u/μL) and 33.75 μL nuclease-free water. The thermocycler program for each PCR reaction began with an initial denaturation step at 95 °C for 2 min followed by 35 cycles of 30 seconds denaturation step at 95 °C, a 30 seconds annealing step at 55 °C, and a 30 seconds elongation step at 72 °C, then one final 5 min extension step at 72 °C.

For the incorporation of biotin molecules into the probe, biotin-16-deoxyuridine (bio-16-dUTP) (Sigma-Aldrich, USA) was substituted for the appropriate unlabeled dNTP in the following polymerase reaction mix. The PCR 2 was carried out with almost the same design as PCR 1 but 1 μL from PCR 1 product was used as template DNA, and in place of 1 μL dNTP mix, 1 μL d (A, C, G) TP (10 mM) and 0.5 μL 16dUTP (0.2 mM) were used. In the course of PCR 2, the biotin 16-dUTP is incorporated into the newly synthesized complementary DNA strands. The final result of this procedure is a 300 bp biotinylated DNA probe with the following sequence: 5’-AACACGCAAATTAATGCCTGTCTGTGTGGAAACTAAAGCCATAG TTTCAACTATACAGCGTAAATATAAGGGTATTAAAATACAAGAGGGTGTGGTTGATTATGGTGCTAGA TTTTACTTTTACACCAGTAAAACAACTGTAGCGTCACTTATCAA CACACTTAACGATCTAAATGAAACTCTTGTTACAATGCCACTTGG CTATGTAACACATGGCTTAAATTTGGAAGAAGCTG CTCGGTATATGAGATCTCTCAAAGTGCCAGCTACAGTTTCTGTTTCTTCACCTGAT GCTGTTAC-3’.

### Fluorescent In situ Hybridization (FISH)

Infected cells were fixed using PFA 4 % pH 7.4 prepared in PBS 1X pH 7.4 DEPC for 20 minutes at room temperature. After that, cells were washed 3X using PBS DEPC solution to remove remains of the fixation process. Following, cells were treated with a 2% H_2_O_2_ solution in methanol for 30 minutes at room temperature and protected from the light, to avoid autofluorescence. Then, cells were washed with PBS DEPC to remove remains of the treatment and incubated for 2 minutes at room temperature in a solution of Proteinase K (10 μg/mL) diluted in PBST 0.1 M pH 7.4 DEPC, to enable cell permeabilization. The reaction was then inactivated using a 2 mg/mL glycine solution, diluted in 0.1M PBST pH 7.4 DEPC for 10 minutes at room temperature. A new fixation was performed using 4% PFA pH 7 for 10 minutes at room temperature. Next, samples were washed with PBS to remove the remains of the fixative.

To a Pre-Hyb step, the Hyb solution (50 % formamide, 10 % dextran sulfate, and 2x SSC pH 7) was used supplemented with salmon sperm DNA (2 μg/mL) to avoid nonspecific signals, for 2 hours at 37 °C in a humid box. Then, the incubation was performed using Hyb solution with the biotinylated probe from PCR 2 (100 μL/mL), denaturing the solution at 85 °C and adding to the cells, incubating for 16 hours at 37 °C in a humid box. To remove Hyb gradually, cells were washed with a 50 % Hyb + 2x SSC pH 7 solution for 20 minutes at 37 °C followed by the repetition of the process using a 25 % Hyb + 2x SSC pH 7 solution. Samples were then washed twice with a 2× SSC solution for 10 minutes at 37 °C, followed by a wash using PBS 1X DEPC for at least 5 times to remove Hyb and SSC remains and probe excess that was not hybridized. Finally, Streptavidin- (Invitrogen #21842) diluted 1:500 in PBS 1× DEPC was used to incubate the samples at room temperature for 2 hours, protected from the light, followed by a washing step using PBS to remove the Streptavidin excess.

DAPI (Santa Cruz Biotechnology - #SC-3598) diluted 1:1000 in PBS solution was incubated for 10 minutes at room temperature and protected from the light. Subsequently, cells were washed with PBS and the labels mounted in an aqueous mounting solution for confocal imaging.

### Immunofluorescence

Cells were prepared onto silanized glass slides, fixed, and stained as previously described in the FISH process. Briefly, after fixation with 4 % PFA, cells were washed with 0.1 M PBST pH 7.4. After, cells were incubated per 10 min with 0.1 M glycine and treated with BSA solution (Sigma) for 30 min. Cells were then incubated overnight at 4 °C with primary antibodies at 1:100 dilution in PBST and 1 % BSA [SARS-COV-2 Spike S1 antibody (#HC2001 GenScript - #A02038), p62 antibody (#BD 610832), Lamp1 antibody (#BD 555798) and CD63 antibody (#BD 556019), according to the desired double IF. The slides were washed and incubated for 2 h with secondary antibodies (Alexa 488 anti-Human IgG Thermo Fisher - #A11013 and Alexa Fluor 555 Anti-Mouse IgG #A21422), diluted 1:500 in PBST+ 1 % BSA. Cells were then washed and stained with DAPI (Santa Cruz Biotechnology, #SC3598). Subsequently, cells were washed with PBS and the labels mounted in an aqueous mounting solution for confocal imaging.

### Confocal microscopy

Microscopic images were acquired with an Airyscan Zeiss LSM880 on an Axio Observer 7 inverted microscope (Carl Zeiss AG, Germany) with a C Plan Apochromat 63x/1.4 Oil DIC objective, 4x optical zoom. Before image analysis, raw.czi files were automatically processed into deconvoluted Airyscan images using Zen Black 2.3 software. DAPI images were acquired as conventional confocal images using a 405 nm laser line for excitation and pinhole set to 1 AU.

### VeroE6 RNAseq

Raw RNA sequencing (RNAseq) reads of VeroE6 cells infected with SARS-CoV-2 at a multiplicity of infection (MOI) of 0.3 for 24 h and mock-infected cells were obtained from SRA (GSE153940) [23,24]. Gene level quantification was performed using Salmon [25] with the *Chlorocebus sabaeus* reference transcriptome obtained from NCBI as the index. Salmon quant mode was run using standard settings and the following flags: --validate mappings and --numBootrstraps 100. The R package tximport [26] was used to load quant.sf files to R and to create a DESeq2 object for differential expression analysis. DESeq2 [27] was used to find differentially expressed genes between SARS-CoV-2 and mock-infected cells using standard settings. Geneset enrichment analysis (GSEA) of differentially expressed genes was performed using the fgsea R package [28] against a custom geneset containing all pathways and terms from Reactome, Gene Ontology (GO) biological process, GO cellular compartment, GO molecular function, Biocarta, KEGG, Hallmark pathways, and Wikipathways. All original GMT files were obtained from the GSEA-MsigDB webservice [29,30]. From the enrichment results containing all terms, pathways related to mTOR signaling, autophagy, lysosome, exocytosis, endosome, late endosome, SNAREs, and vesicle-associated proteins were selected for visualization. Only terms with an adjusted p-value < 0.05 were considered for discussion. Leading-edge genes from the significant terms were used to depict results at the gene level. Figures were elaborated using the ggplot2 and pheatmap R packages.

### Single-cell RNAseq (scRNAseq) of severe COVID-19 patients

Single-cell transcriptomic data was analyzed from bronchoalveolar lavage fluid (BALF) from patients with varying severity of COVID-19 and matching healthy controls [31]. The dataset generated by the original authors is publicly available at https://covid19-balf.cells.ucsc.edu/. Briefly, the dataset was downloaded and the RDS seurat object was imported into R environment version v3.6.3. Epithelial cells were selected for downstream analysis based on the cell type annotation provided by the authors. Cells containing viral reads according to the original author’s annotation were labeled as “infected” and those in which no viral reads were detected were labeled as “bystanders”. Only cells from severe COVID-19 patients presented viral reads, therefore cells from moderate patients were discarded. Differential expression analysis was conducted using the FindMarkers function of the seurat package v3.1 [32] using Wilcoxon test to compare infected and bystander cells of severe patients to healthy control cells and also infected to bystander cells of severe patients. Differentially expressed genes were identified considering genes expressed in at least 10% of cells, FDR < 0.05 and | avg_logFC| > 0.25. Differentially expressed genes found in each individual comparison were submitted to GSEA using the fgsea R package with the same custom geneset file and selected terms used for the VeroE6 analysis described above. Violin plots of the expression of selected differentially expressed genes were produced using the VlnPlot function of the seurat package.

### Western blotting

Proteins were separated by SDS-PAGE and transferred onto nitrocellulose membranes. Nitrocellulose membranes were blocked in a solution of TBS containing 5 % nonfat dry milk and 0.1 % Tween-20 for 2 h with constant agitation. After blocking, the membranes were incubated with anti-mTOR (Cell Signaling, #2972), anti p-mTOR (Cell Signaling, #2971), p-S6K1 (Cell Signaling, #9234), anti-S6K1 (Cell Signaling, #2708), anti-pS6 (Cell Signaling, #2215), anti-S6 (Cell Signaling, #2317), anti-pULK1 (Cell Signaling, #5869), anti-ATG7 (Cell Signaling, #8558), anti-LC3 (Cell Signaling, #4108), anti-p62 (Cell Signaling, #88588), and anti-β-actin (Cell Signaling, #4967), antibodies overnight at 4 °C. Membranes were washed with TBS-T (3 times for 10 min) and incubated with horseradish peroxidase-conjugated secondary antibodies anti-mouse (Millipore, #AP308P), or anti-rabbit (Thermo Scientific, #31460) according to the primary antibody for 1 h at room temperature with constant agitation. Bands were visualized using the ECL kit (GE Healthcare). Band densitometry was measured using ImageJ software.

### Quantification and Statistical Analysis

The software packages GraphPad Prism 8.0 and 9.0 were used for statistical analyses of Western blotting and PFU, respectively. The values presented are means and standard deviation (SD). The mean difference was tested by Test T analysis. p<0.05 were considered significant.

## 3. Results

### 3.1. SARS-CoV-2 infection modulates mTOR signaling and autophagy in Vero E6 cells

We analyzed the mTOR and autophagy pathways in SARS-CoV-2 infected Vero E6 cells, measuring the levels of mTOR, S6K1 (Ribosomal protein S6 kinase beta-1), and S6 (Ribosomal protein S6) phosphorylation and LC3I/LC3II (Microtubule-associated protein light chain 3) and p62/SQSTM1 (Ubiquitin-binding protein p62/Sequestosome 1) protein content (Figure 1 and Figure S1). SARS-CoV-2 infected cells exhibited increased levels of mTOR, S6K1, and S6 phosphorylation by over 3-fold, compared to the baseline levels of mock cells (Figure 1A). The significantly increased mTOR and S6 phosphorylation levels can also be observed when we use the endogenous gene for their quantification (Figure S2). SQSTM1/p62 levels and LC3 conversion usually correlate with autophagic flux [15,22].

**Figure 1.**
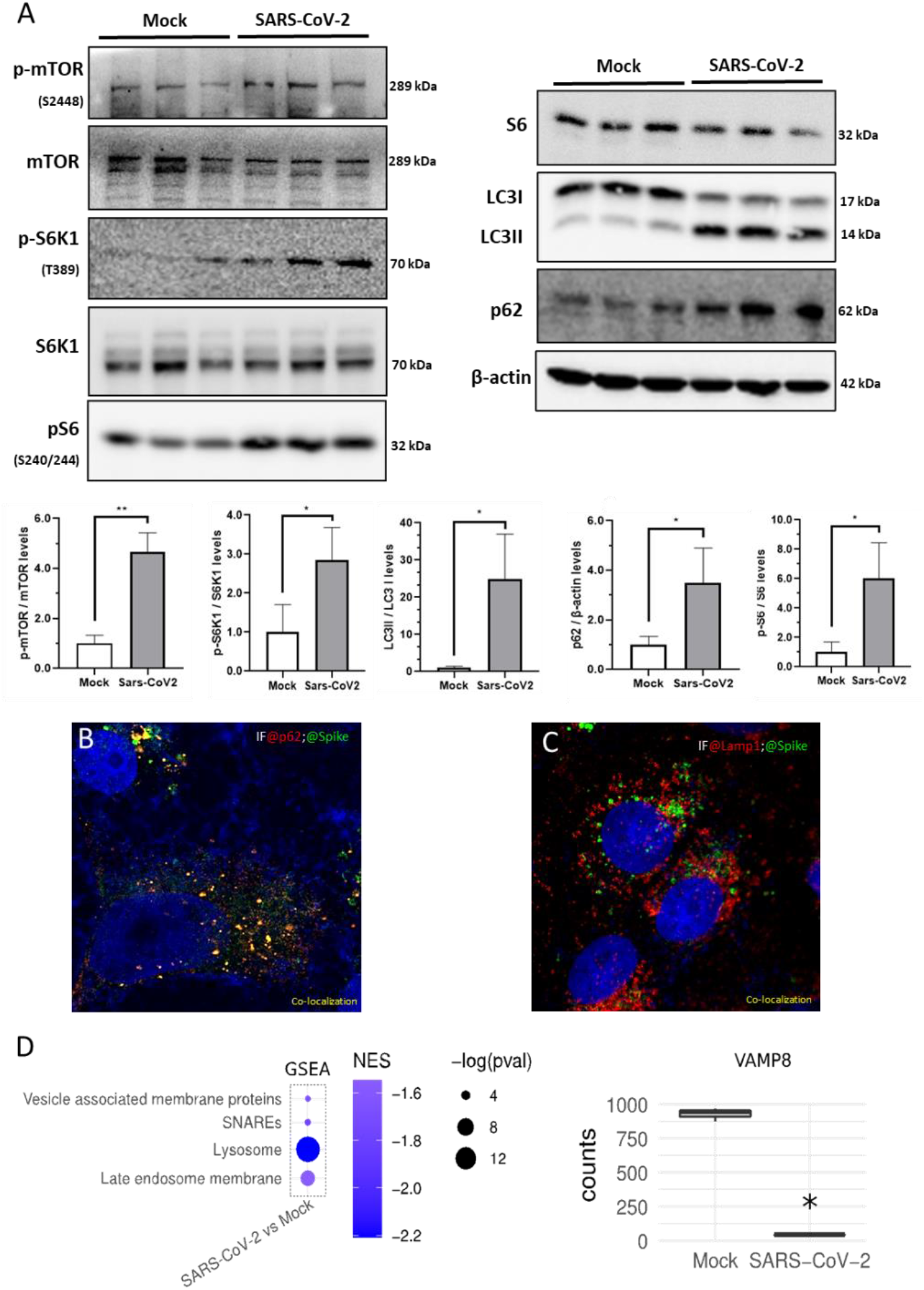
SARS-CoV-2 infection activates the mTOR pathway and impairs autophagy in Vero E6 cells. **(A)** mTOR, p-mTOR, S6K1, p-S6K1, S6, p-S6, LC3I/II, p62, and N immunoblotting of Vero E6 cells protein extracts 24 hpi. β-actin and Vinculin were used as endogenous control. Raw images are available in Figure S2. **(B)** IF demonstrating co-localization (yellow dots) of autophagosome protein p62 and SARS-CoV-2 Spike protein in Vero E6 cells M.O.I.=1. **(C)** IF for lysosomal protein Lamp1 (red) and SARS-CoV-2 Spike protein (green) showing no co-localization (yellow) in Vero E6 cells MOI=1. **(D)** Downregulation of genes related to lysosomal activity and of VAMP8, a SNARE responsible for autophagosome-lysosome fusion, in Vero E6 cells. All experiments were infected with MOI =1 Data represent means ± SD in samples from independent experiments in each experiment. T-test was used for immunoblotting analysis. *p<0.05 were considered statistically significant.

Using confocal microscopy, immunofluorescence (IF) for p62 and the viral Spike protein revealed that SARS-CoV-2 colocalizes within autophagosomes in infected cells (Figure 1B), but not with Lamp1 staining (Figure 1C). RNAseq data from SARS-CoV-2-infected Vero E6 cells [23] revealed the downregulation of genes related to lysosomal activity and of the VAMP8 (Vesicle-associated membrane protein 8). VAMP8 is a SNARE (soluble N-ethylmaleimide-sensitive factor attachment protein receptors) involved in autophagosome-lysosome fusion [33,34] (Figure 1D).

These results demonstrate that SARS-CoV-2 infection in Vero E6 cells induces accumulation of autophagosome proteins in a cell environment with increased mTOR activity, a condition known to inversely correlate with lysosomal activity [35,36]. At the same time, the virus infection is associated with the repression of genes related to lysosome and vesicle fusion.

### 3.2. Severe COVID-19 patients display hyperactivation of mTOR signaling and suppression of autophagic flux and VAMP8

We then asked whether COVID-19 patients samples would also present some of the alterations observed in SARS-CoV-2 infected Vero E6 cells. We analyzed a publicly available single-cell RNAseq (scRNAseq) dataset of epithelial cells derived from bronchoalveolar lavage fluid (BALF) of severe COVID-19 patients [37] (Figure 2A). scRNAseq allows the detection of the virus RNA in individual cells and the comparison of genome-wide gene expression changes between the infected cells and bystander (not infected) cells and cells of healthy controls. We found that both infected and bystander BALF epithelial cells of severe COVID-19 patients presented upregulation of mTOR signaling, repression of autophagy genes, and upregulation of negative regulators of autophagy, all compared to healthy controls (Figure 2B-C). The same was shown for Vero E6 cells (Figure 1A and D), as already described. The expression of genes related to mTOR signaling was not altered in infected cells relative to bystanders (Figure 2B). We suggest either a pre-existing condition of high mTOR signaling and compromised autophagy flux in severe COVID-19 patients or a cell effect elicited by the virus.

**Figure 2.**
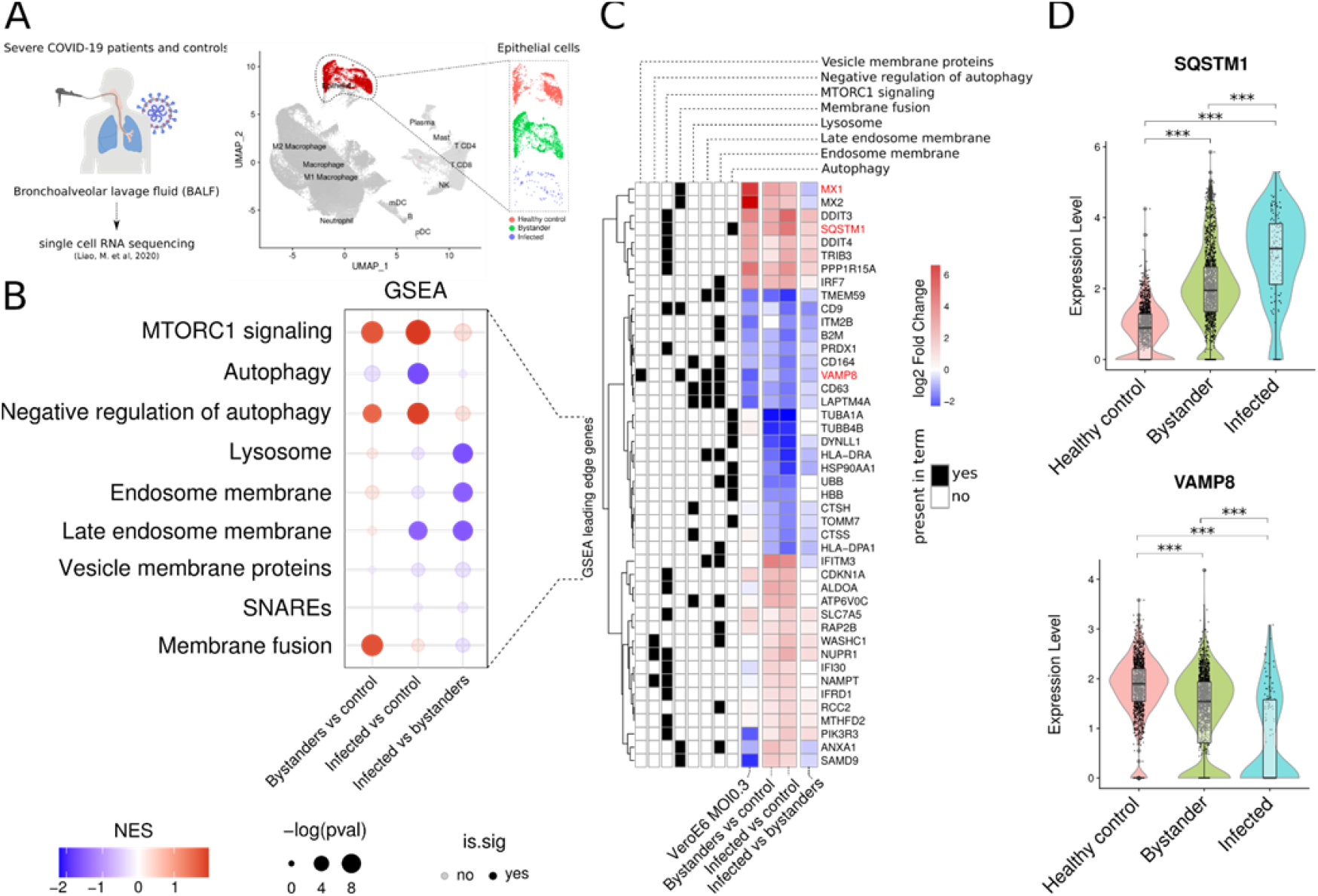
scRNAseq analysis in COVID-19 patients. **(A)** Bronchoalveolar lavage fluid (BALF) samples of severe COVID-19 patients and matched controls were submitted to scRNAseq [37]. Epithelial cells were isolated from the other cell types and those in which viral reads were detected were labeled as infected. The remaining cells of COVID-19 patients were labeled as bystanders. **(B)** Gene Set enrichment analysis (GSEA) of differentially expressed genes in severe COVID-19 patient bystander or infected BALF epithelial cells versus control cells or infected versus bystander cells of severe patients. **(C)** Fold change values (log2FC) of the leading-edge genes of enriched Reactome terms depicted in B for SARS-CoV-2 infected VeroE6 cells and scRNAseq of severe COVID-19 patients BALF epithelial cells. **(D)** Expression level of SQSTM1 and VAMP8 in control (red), bystander (green) and infected (blue) BALF epithelial cells. *** differential expression adjusted p-value < 0.01.

Differential gene expression analysis between infected and bystander BALF epithelial cells of severe COVID-19 patients recapitulated the downregulation of genes related to lysosome and vesicle membrane proteins (Figure 2B-C) observed in Vero E6 cells. *VAMP8* was consistently downregulated in both infected and bystander BALF epithelial cells, with the lowest level of expression observed in infected cells from severe COVID-19 patients (Figure 2D). *SQSTM1*, the gene encoding the p62 protein, on the other hand, was upregulated in bystander or infected cells from severe COVID-19 patients compared to healthy controls, with the highest expression detected in infected cells (Figure 2D).

### 3.3. Vero E6 cells accumulate SARS-CoV-2 particles inside cells

In addition to the role of VAMP8 in autophagosome-lysosome fusion, this protein is also necessary for endosome and secretory vesicle fusion with the plasma membrane [38–40]. Since downregulation of VAMP8 and upregulation of mTOR genes involved in membrane fusion were demonstrated in both SARS-CoV-2 infected Vero E6 cells and BALF epithelial cells of severe COVID-19 patients, we next explored the possibility that SARS-CoV-2 infection could block exocytosis and reduce viral release. We compared the amount of viable SARS-CoV-2 particles (plaque-forming units - PFU) inside Vero E6 cells to the number of PFUs in the extracellular medium 24 hours post-infection (hpi) (Figure 3). 20 times more SARS-CoV-2 particles were detected inside the cells than on the supernatant at 8 hpi, and 25 times more viral particles were retained inside the cells at 24 hpi (Figure 3A).

**Figure 3.**
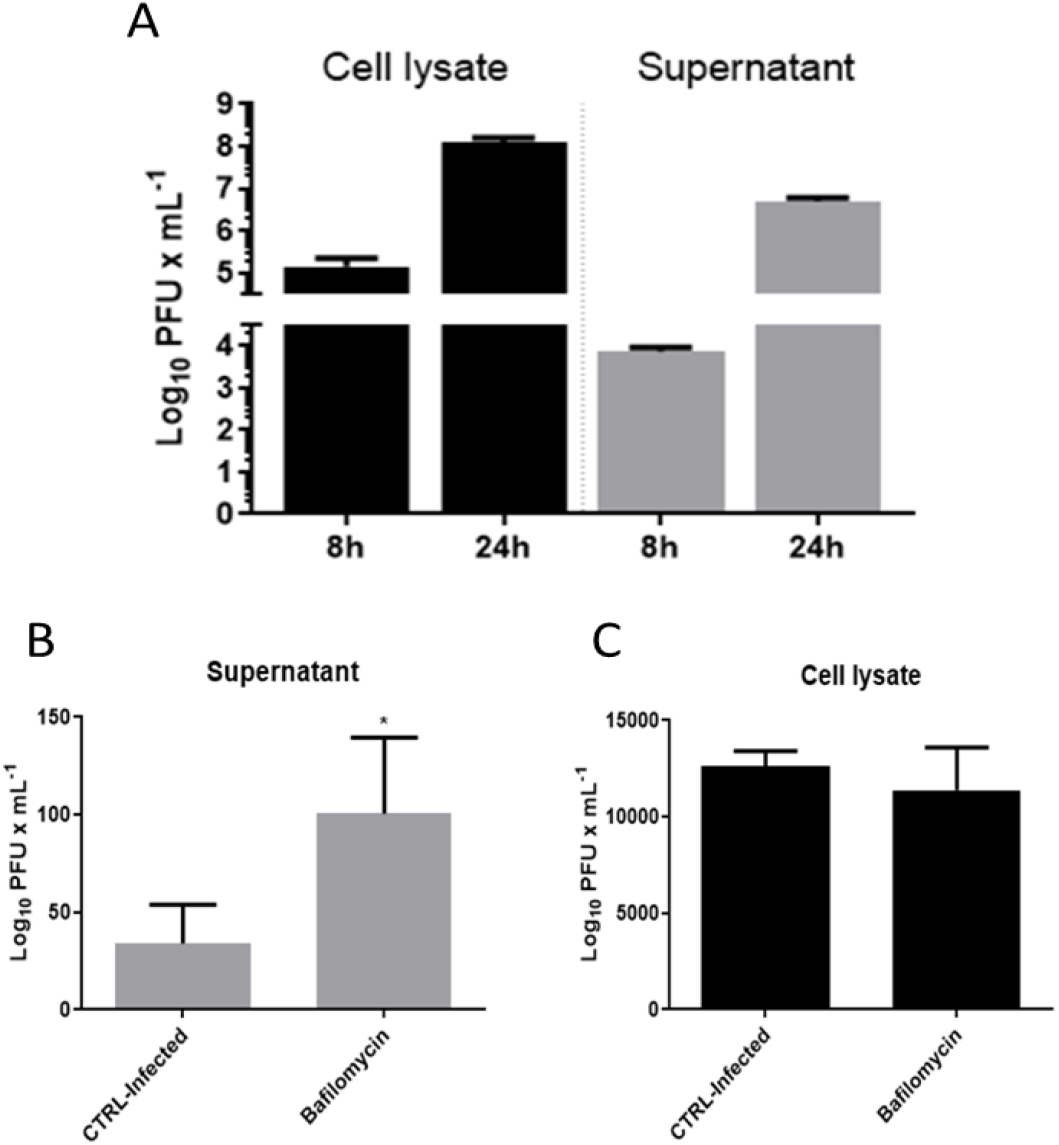
PFU and Bafilomycin treatment 24 hpi in Vero E6 cells. **(A)** Plaque-Forming units assay of cell lysate and supernatant of Vero E6 cell culture 8 and 24 hpi with SARS-CoV-2. **(B)** Plaque-Forming units assay of cell lysate and **(C)** supernatant of Vero E6 cell culture treated with 10 nM bafilomycin after 24 hpi with SARS-CoV-2. Bars represent mean ± SEM. Statistic difference is indicated by *p<0.05.

These findings, combined with the repression of genes related to exocytosis in SARS-CoV-2 infection (Figure 2C-D), suggest that the retention of viral particles inside infected cells could result from hampered exocytosis. To test this hypothesis, we treated SARS-CoV-2 infected Vero E6 cells with the exocytosis-inducer bafilomycin [22,41]. Bafilomycin A1 is an inhibitor of vacuolarH+-ATPase (vATPase), an enzyme essential for lysosome acidification and degradation of autophagosome cargo [14]. The bafilomycin treatment did not significantly change the amount of SARS-CoV-2 particles inside the cells, but increased viral particles in the supernatant of bafilomycin-treated cells (Figure 3B-C). These results suggest that exocytosis inhibition limits viral release to the extracellular medium in SARS-CoV-2 infected cell culture.

### 3.4. SARS-CoV-2 cell-to-cell transmission in CD63-positive vesicles

We then performed fluorescent in situ hybridization (FISH) and double IF to study the dynamics of newly formed SARS-CoV-2 particles inside the cells. In addition to the dot-like staining co-localized with p62, we identified vesicle-like labeling of the virus genetic material (Figure 4A) and viral proteins (Figure 4B), that co-localized entirely with CD63 (Figure 4C-D), a marker for late endosome/multivesicular body (MVB). Strikingly, these virus-bearing CD63-positive vesicles were found in cell-to-cell connections at 24 hpi (Figure 4D). These observations suggest that, upon assembly, viral particles loaded into CD63-positive vesicles accumulated inside infected cells, which may be transported to neighboring cells via cellular connections.

**Figure 4.**
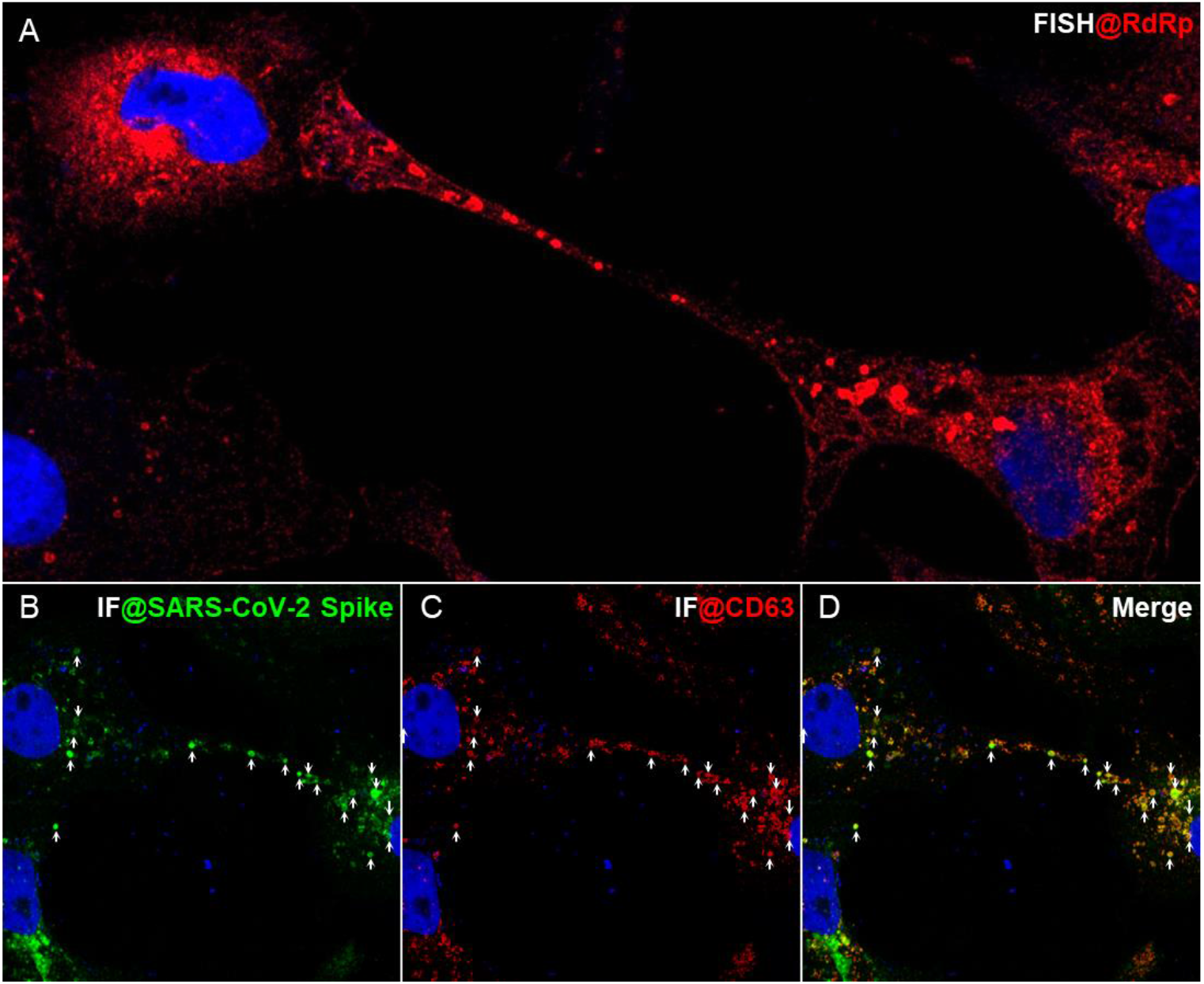
SARS-CoV-2 detection in cell-to-cell connections in Vero E6 cells by fluorescent in situ hybridization. **(A)** and immunofluorescence **(B, C, D).** FISH for the RNA-dependent RNA polymerase gene showing vesicle-like labeling in a connection between two neighboring cells **(A)**. IF for Spike protein **(B)** double-stained with CD63 **(C)**, demonstrating viral-containing late endosomes/MVBs in cell-to-cell connections **(D)**.

## 4. Discussion

This study reports that SARS-CoV-2 infection induces transcriptional activation of autophagosome proteins and represses genes related to lysosomes and vesicle fusion in a scenario with high mTOR activity and repressed autophagy. We demonstrated these effects both in Vero E6 cells and in COVID-19 patients samples. By studying SARS-CoV-2 infection *in vitro*, using the Vero E6 cell line as a model, we further demonstrated that newly assembled viral particles are localized to autophagosomes and MVBs/late endosomes. Both organelles can be targeted to lysosomal degradation but, instead, they seem to accumulate in SARS-CoV-2 infected cells. We also observed SARS-CoV-2 viral particle retention in Vero E6 cells 24 hpi and demonstrated that CD63-positive vesicles loaded with viral particles are found in cell-to-cell connections (Figure 4D).

The findings described in this report have major implications for our understanding of the SARS-CoV-2 infection and the causal factors for COVID-19 severity. First, although we found activation of the mTOR signaling in SARS-CoV-2 infected cells, our single-cell analysis of severe COVID-19 patients revealed that mTOR signaling is not altered between infected and bystander cells. However, this effect is detected by comparing bystander or infected cells to healthy controls. The same is observed for autophagy-related genes, reduced between COVID-19 patient cells and healthy controls, but not between infected and bystander patient cells. The results suggest that high mTOR signaling and compromised autophagy flux could be a cell response to the viral infection or pre-existing conditions that could contribute to the poor prognosis of the disease. In addition to blocking autophagy flux, high mTOR activity could favor cap-dependent translation of SARS-CoV-2 genomic and 3’-co-lateral capped subgenomic mRNAs [42–45].

mTOR signaling has been proposed as a factor linking obesity to COVID-19 severity [3] and was demonstrated to be modulated by SARS-CoV-2 infection in Huh7 [46] and Vero E6 [2,47] cell lines (Figure 1). In line with a possible causal role of high mTOR signaling favoring COVID-19 severity, mTOR inhibition resulted in a significant reduction in virus replication and may also inhibit viral particle uptake in Huh7, Vero E6 cells and mucociliary primary human airway derived air-liquid interface cultures [46,47]. Hyperactivation of mTOR signaling is present in hypertension [48], type-2 diabetes [49], including related to aging [50], ischemic heart disease [51], atrial fibrillation [52], chronic renal failure [53] dementia caused by Huntington’s [54] and Alzheimer’s [55] diseases, Chronic Obstructive Pulmonary Disease (COPD) [56], cancer [57], heart failure [58] and obesity [59]. These are ten comorbidities considered most prevalent in COVID-19 deceased patients (Palmieri et al., 2020). It is also known that elders are the most vulnerable group when affected by COVID-19, corresponding to 80% of hospitalized patients [1]. Increased mTOR signaling and reduced autophagy are hallmarks of aging [50]. Interventions that reduce mTOR signaling and promote autophagy prolong lifespan in different species and protect from diseases, including infectious diseases [60,61]. Indeed, the inhibition of mTOR pathway by mTORC1 inhibitors such as rapamycin and rapalogs reduced SARS-CoV-2 infection and N protein expression in Vero E6 and primary human airway-derived air-liquid interface cultures, endorsing our results [47].

Another important aspect of our findings is the dependence of SARS-CoV-2 on autophagosomes and the viral-induced blockage of the autophagic flux. We demonstrated here that SARS-CoV-2 colocalizes within autophagosomes in infected, but not with Lamp1. These results corroborate previous evidence that autophagosome-lysosome fusion is blocked during SARS-CoV-2 infection [62]. Several other groups have shown that coronaviruses can increase autophagosome accumulation and block their fusion with lysosomes through direct interactions between viral and cellular proteins [14,15,62,63]. As demonstrated here for SARS-CoV-2, RNA viruses, such as the Hepatitis C Virus (HCV) and the Human Parainfluenza Virus Type 3 (HPIV3), hijack the autophagic machinery and lead to the accumulation of LC3II and p62 [64–67]. These strategies improve viral replication and improve the ability of these viruses to evade the host’s immune system [66,68–72]. For HPIV3, the impairment of the autophagic flux is caused by a virus-induced disruption of the function of the SNARE protein Synaptosome Associated Protein 29 (SNAP29), blocking its interaction with Syntaxin17 (STX17) and the formation of the SNAP-STX17-VAMP8 complex [65].

Although we did not find significant differences in the expression of the SNAP29 or STX17 genes between infected cells and controls in our analyses (data not shown), we demonstrate that VAMP8 is drastically repressed in infected Vero E6 cells and the BALF epithelial cells of severe COVID-19 patients. Recently, a study showed that overexpression of SARS-CoV-2 ORF3a protein in Vero E6 cells blocks the STX17-SNAP29-VAMP8 SNARE complex assembly, impairing the autophagic flux and late endosome integrity in infected cells. However, ORF3a overexpression alone did not significantly affect VAMP8 gene expression [62]. Therefore, the consistent repression of VAMP8 in Vero E6 cells and severe COVID-19 patients samples presented here cells also suggests that SARS-CoV-2 affects the SNARE complex also at the transcriptional level. Interestingly, increased mTOR activity is also essential for suppressing lysosomal biogenesis, autophagy, and lysosomal acidification [73], which might also be required to facilitate the fusion between autophagosomes and lysosomes [74,75]. These observations also support the role of increased mTOR signaling in COVID-19 severity.

A recent multi-omics study [76] revealed that SARS-CoV-2 infection induces phosphorylation of key proteins in the mTOR pathway, including SQSTM1. The study also shows that SARS-CoV-2 infection promotes ubiquitination of autophagy-related proteins, including VAMP8, suggesting protein degradation via the proteasome. The results of this multi-omics study are in agreement with our findings, showing that SARS-CoV-2 regulates mTOR activity and autophagy at different levels. Nonetheless, another study showed that SQSTM1/p62 levels did not change during SARS-CoV-2 infection [21]. We demonstrated that the ratio of the autophagosome markers LC3-II to LC3-I and the content of p62/SQSTM1 increase in infected cells (Figure 1A), which is associate with the accumulation of autophagosomes by blockage of the autophagy flux [22].

SARS-CoV-2 infection in Vero E6 cells also resulted in the reduction of virus release from infected cells and transport of virus-loaded CD63-positive vesicles between neighboring cells (Figures 3 and 4). In contrast, SARS-CoV, which caused the 2003 SARS epidemics, presents ten times more viral particles in the supernatant than inside the cells at 12 hpi in Vero E6 cells [77]. MVBs fuse with the plasma membrane during exocytosis and release their intraluminal vesicles to the extracellular space as exosomes [78,79]. Although the exact mechanisms preventing SARS-CoV-2 release from cells need further clarification, we found that the expression of genes related to vesicle membrane and fusion, such as VAMP8 [33], are robustly repressed in Vero E6 infected cells and BALF epithelial cells of severe COVID-19 patient. Once again, elevated mTOR signaling found in COVID-19 patients might be sufficient to prevent exocytosis. The transcription factor EB (TFEB), which mTORC1 negatively regulates, is necessary for exocytosis of lysosomes [80] and autophagolysosomes [81]. This regulation occurs through the regulation of SNARE proteins syntaxins 4 and 6, which are necessary for the fusion of transport vesicles with the plasma membrane [82].

While higher resolution details and dynamics would help to dissect how vesicular traffic is affected by SARS-CoV-2 infection, we demonstrate here that virus-loaded CD63-positive vesicles accumulate inside the cells and are observed in membrane bridges between neighboring cells. SARS-CoV-2 viral particles have been observed in cellular protrusions in Caco-2 cells [83] and tunneling-nanotubes (TnT) between neighboring cells in Vero E6 cells. However, unlike what is shown here (Figure 4), the viral particles were detected at the outer surface of these membrane bridges connecting Vero E6 cells [84]. The cell-to-cell transfer of viral components demonstrated here might represent an additional mechanism that SARS-CoV-2 uses to infect new cells while escaping the immune system and shed light on the virulence mechanisms and transmissibility SARS-CoV-2. For example, viral particle accumulation inside cells could explain cases of persistent systemic infection by SARS-CoV-2, the late onset of the symptoms, and the fact that many asymptomatic individuals are still able to produce infectious viral particles [21,85,86].

This study was experimentally performed using Vero E6 cell culture, a largely accepted cell model for SARS-CoV-2 infection [87]. This model is limited since Vero E6 cells are derived from a non-human primate and are interferon-deficient [88]. Several in vitro studies performed in Vero E6 and human cell lines suggested promising effects of drugs on SARS-CoV-2 infection in cell lines [89–93] but few have been translated to the infection in patients. For this reason, we validated our hypothesis using a single-cell RNAseq dataset of severe COVID-19 patient lung epithelial cells. We detected similar gene expression results between the VeroE6 and patient experiments, supporting that SARS-CoV-2 blocks the autophagic flux and reduces vesicle membrane fusion, likely reducing viral release from the host cells and relating to a cell-to-cell viral transmission. However, it is still necessary to confirm that this phenomenon also occurs in human cells and tissues and is clinically relevant in COVID-19 patients.

Our findings may support the design new or the interpretation of current clinical trials of FDA-approved mTOR inhibitors. There are several active clinical trials registered on ClinicalTrials.gov for mTOR inhibitors, such as the rapamycin analog Sirolinus. These clinical trials are based on the proposed effects of rapamycin on restoring T-cell functionality and decrease cytokine storm [94,95], the direct antiviral effect [96], and in aging-related declining of the immune function [97]. Metformin, a medicine commonly used to treat Type 2 Diabetes and a well-known mTOR inhibitor [98], has been demonstrated to reduce COVID-19 mortality [99]. Among the possible mechanisms proposed to explain metformin’s positive effect on COVID-19 outcomes, the inhibition of the mTOR pathway could reduce the severity of COVID-19 disease by preventing immune hyperactivation [100,101].

Based on findings presented here we constructed the model presented in Figure 5. conditions of low mTOR signaling, autophagy is promoted, and vesicles containing SARS-CoV-2 particles might fuse to acidified lysosomes and be eliminated or go directed to exocytosis, exposing the virus to the immune system (Figure 5A). The SARS-CoV-2 infection seems to be favored by high mTOR activity, which could guarantee cap-dependent translation of viral proteins and lysosome deacidification, allowing viral assembly in autophagosomes and late endosomes (Figure 5B). Therefore, the machinery that promotes SARS-CoV-2 viral replication appears to prevent viral release, resulting in the accumulation of novel viral particles inside the cells, which may contribute to cell-to-cell transmission and evasion of the immune system.

**Figure 5.**
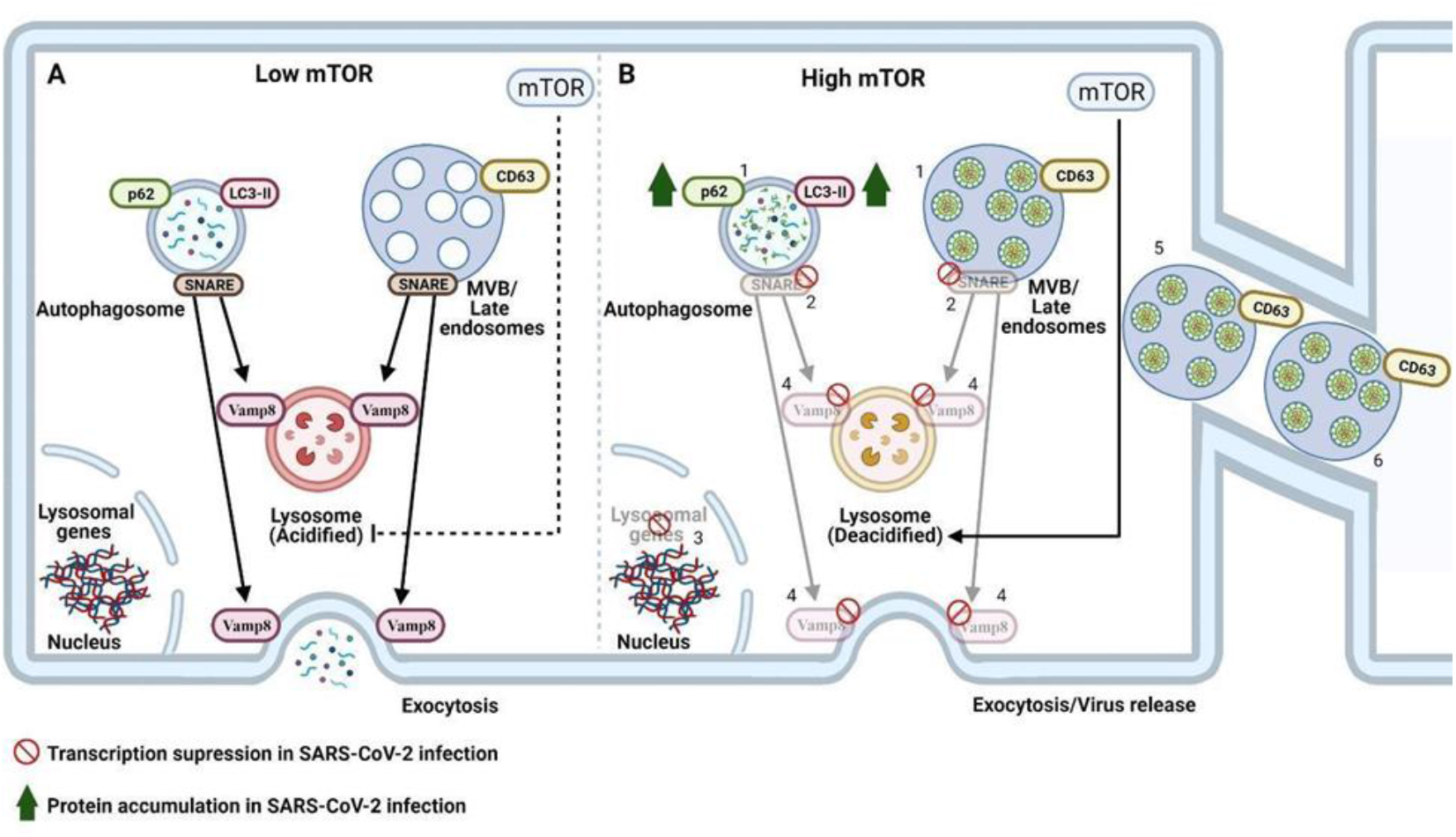
Proposed molecular mechanisms of SARS-CoV-2 infection upon high mTOR activation and compromised autophagy flux and vesicle fusion. **(A)** Classical autophagy flux, in low mTOR conditions, where SNARE proteins allow for vesicle fusion to acidified lysosome and vesicles fusion to the cell membrane. **(B)** High mTOR scenario depicting a SARS-CoV-2 viral infection. Newly assembled viral particles, associated with cellular vesicles (1), are prevented from being destroyed by lysosomal fusion as the expression of genes coding for vesicle fusion proteins (2) and lysosomal genes (3) are down-regulated, lysosomes are deacidified (31), and lysosomal fusion to virus-bearing vesicles, prevented (31, 32). The suppression of vesicle-membrane fusion proteins, such as VAMP8 (4), would prevent virus release from infected cells. Virus-bearing MVBs accumulate in infected cells (5) and are observed in cell-to-cell connections (6). Labeling 1-6 represents the original evidence of this manuscript. Created with BioRender.com

## 5. Conclusions

Here we show that mTOR activity and impairment of autophagy are important events of SARS-CoV-2 infection in vitro and in COVID-19 patients samples. Besides, markers of lysosome acidification like VAMP8 were demonstrated to be downregulated by SARS-CoV-2 infection, impacting the dynamics of viral particles released from cells. Finally, the subcellular localization of virus components presented here indicates complex mechanisms of cell-to-cell transmission, that are still to be further explored. In sum, we present evidence that the mTOR and autophagy pathways are important players in the dynamics of SARS-CoV-2 in the cell.

## Supplementary Materials

The following are available online at www.mdpi.com/xxx/s1, **Figure S1**: SARS-CoV-2 infection activates the mTOR pathway and impairs autophagy in Vero E6 cells./ **Figure S2:** Raw membranes of all immunoblotting assays

## Author Contributions

GCM, TLD, BB, MRA and EPZ led and performed experiments, literature search and data plotting and analyses. LNS; LB; JGAE; CM performed IF and HIS. MCSM, ICBP, APM, EPZ and LGSS performed WB. DATT, KBS, PLP, SPM, GFS and FG performed all SARS-CoV-2 infections in Vero E6 cell and PFU. IC performed bioinformatics. GOB performed confocal image processing. RGL, TLK and TDS performed cellular experiments. HC and LLPS contributed with reagents, insights and interpretation and discussions of the results. TLD, GCM, MRA and HMS generated all figures. GCM, MCSM, ICBP, FMS and HMS conceived the illustrative scheme. GCM and HMS formatted the manuscript. HMS, FMS, JLPM, HN and MAM supervised the study, interpreted the results and contributed to the discussion. HMS and FMS conceived and designed the study and wrote the manuscript. All the authors revised the manuscript and approved the final version for publication.

## Funding

This work was supported by grants from FAEPEX-UNICAMP (2005/20; 2319/20; 2432/20; 2274/20), São Paulo Research Foundation (FAPESP) (18/14933-2; 20/05284-0; 2014/50938-8; 2020/05346-6; 2020/04919-2; 2020/04558-0) and National Council for Scientific and Technological Development (CNPq) (465699/2014-6).

## Acknowledgments

The authors acknowledge the National Institute of Science and Technology of Photonics Applied to Cell Biology (INFABIC) and the technical support of Mariana Ozello Baratti, for aid with confocal microscopy and Elzira Saviani and Thais Theizen for technical support.

## Conflicts of Interest

The authors declare no conflict of interest.

## Notes

### Competing Interest Statement

The authors have declared no competing interest.

