## Supplementary Figures for "Increased mTOR signaling, impaired autophagic flux and cell-to-cell viral transmission are hallmarks of SARS-CoV-2 infection"

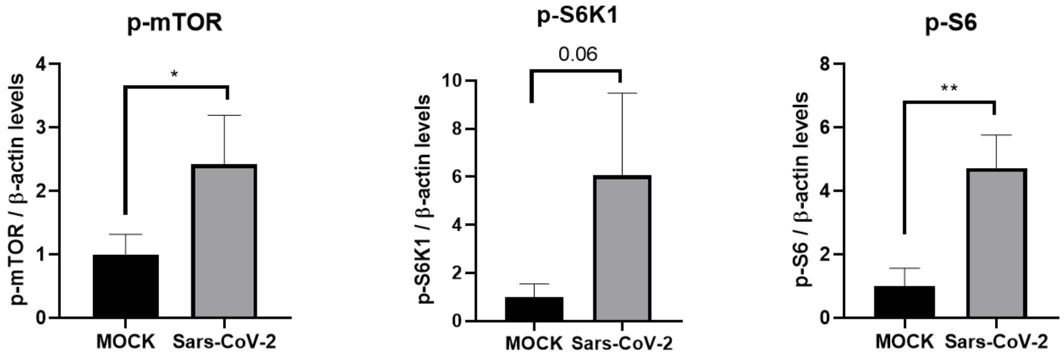


**Figure S1. SARS-CoV-2 infection activates the mTOR pathway and impairs autophagy in Vero E6 cells.** Quantification of p-mTOR, p-S6K1, and p-S6 immunoblotting of Vero E6 cells protein extracts 24 hpi by endogenous β-actin. All experiments were infected with MOI =1 Data represent means ± SD in samples from independent experiments in each experiment. T test was used for immunoblotting analysis. *p<0.05 were considered statistically significant.

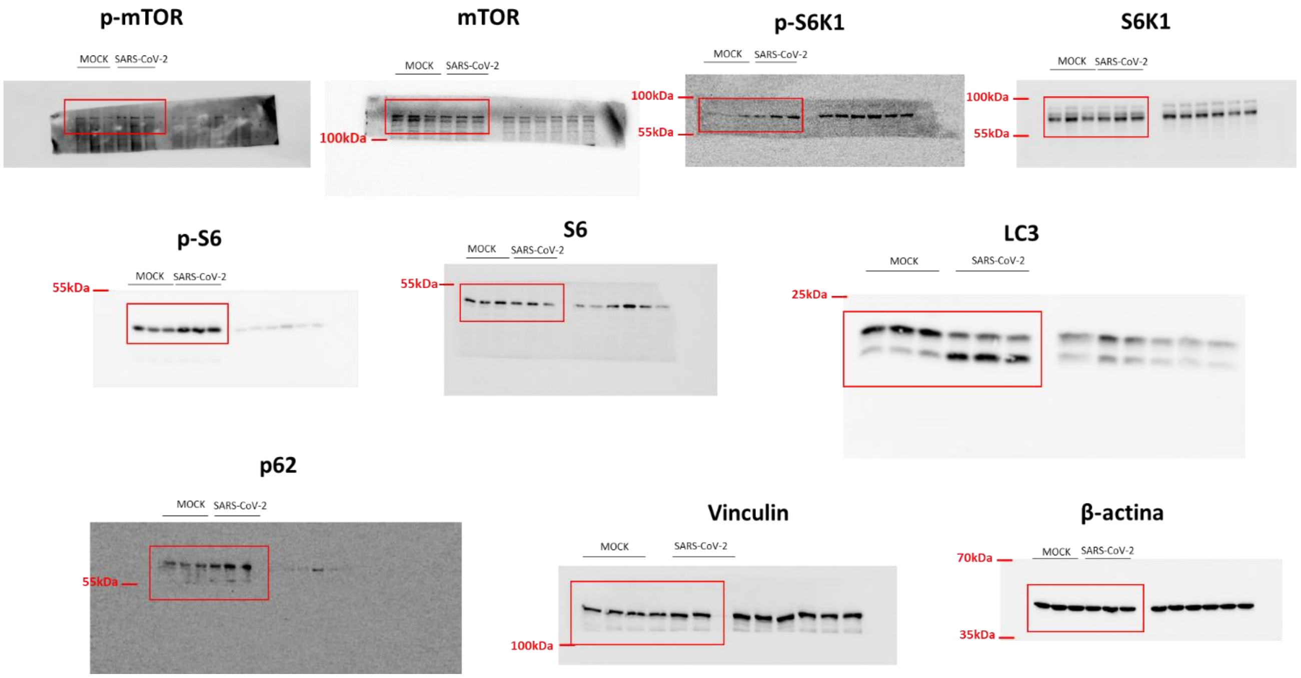


**Figure S2. Raw membranes of all immunoblotting assays.**
